# Shank3 influences mammalian sleep development

**DOI:** 10.1101/2021.03.10.434728

**Authors:** Elizabeth Medina, Hannah Schoch, Taylor Wintler, Kaitlyn Ford, Kristan G. Singletary, Lucia Peixoto

**Affiliations:** Department of Translational Medicine and Physiology. Sleep and Performance Research Center. Elson S. Floyd College of Medicine, Washington State University, Spokane, Washington, United States

**Keywords:** Autism Spectrum Disorder, mouse model, sleep, development, Shank3, EEG, REM, sleep latency

## Abstract

Sleep problems are prevalent in Autism Spectrum Disorder (ASD), can be observed before diagnosis, and are associated with increased restricted and repetitive behaviors. Therefore, sleep abnormalities may be a core feature of the disorder, but the developmental trajectory remains unknown. Animal models provide a unique opportunity to understand sleep ontogenesis in ASD. Previously we showed that adult mice with a truncation in the high-confidence ASD gene *Shank3* (Shank3^ΔC^) recapitulate the clinical sleep phenotype. In this study we used longitudinal electro-encephalographic (EEG) recordings to define, for the first time, changes in sleep from weaning to young adulthood in an ASD mouse model. We show that Shank3^ΔC^ mice sleep less overall throughout their lifespan, have increased rapid eye movement (REM) sleep early in life despite significantly reduced non-rapid eye movement (NREM) sleep, and have abnormal responses to increased sleep pressure that emerge during a specific developmental period. We demonstrate that the ability to fall asleep quickly in response to sleep loss develops normally between 24 and 30 days in mice. However, mutants are unable to reduce sleep latency after periods of prolonged waking and maintain the same response to sleep loss regardless of age. This phenomenon seems independent of homeostatic NREM sleep slow-wave dynamics. Overall, our study recapitulates both preclinical models and clinical studies showing that reduced sleep is consistently associated with ASD and suggests that problems falling asleep may reflect abnormal development of sleep and arousal mechanisms.

**SIGNIFICANCE STATEMENT:** In this first longitudinal sleep study in an Autism mouse model, we demonstrate that sleeping less seems a core feature of the disorder while problems falling asleep emerge during development.

## INTRODUCTION

Good sleep is a cornerstone for maintaining optimal health. Accordingly, sleep problems, with insomnia as the most common sleep disorder, impact significantly on our physical capabilities and mental health. In individuals diagnosed with Autism Spectrum Disorder (ASD), two thirds have chronic insomnia (defined as persistent problems falling and staying asleep), and 86% of people with ASD are affected by sleep problems^1,2^. Relative to individuals that do not have ASD, individuals with ASD experience significant delays falling asleep, multiple night awakenings, and overall less sleep time^3^. Poor sleep is predictive of the severity of ASD core behavioral diagnostic symptoms such as social skill deficits and stereotyped behavior. With age, sleep problems often worsen and heavily affect the quality of life of individuals and their caregivers. In addition, sleep problems in young children that go on to be diagnosed with ASD are associated with increased ‘higher-order’ restricted and repetitive behaviors later in childhood^4^ and altered patterns of brain development^5^. Although commonly referred to as a condition co-morbid with ASD, when children are diagnosed with ASD, they often already have a history of sleep problems, sometimes starting as early as the first year of life. Given the well-documented role of sleep in brain development, early-life sleep disruption is likely to contribute to later-life core features of ASD and might even be an indication for early intervention. However, the developmental trajectory of sleep problems in ASD and its potential role in ASD etiology remains largely unexplored.

ASD is known to have a strong genetic component including both de novo and inherited gene variations. Nonetheless the same variant can cause different symptoms along a spectrum^6^. This heterogeneity presents challenges for genetic ASD animal models in which targeting a single gene of interest yields inconsistent expression of core behavioral phenotypes such as social communication deficits and stereotyped behaviors. In addition, this approach generally fails to capture earlier neurodevelopmental processes preceding disease onset, leaving etiology elusive. Sleep, unlike many other behavioral phenotypes, can be objectively quantified in mammals and is very highly conserved across the animal kingdom. Therefore, animal models are ideally suited to investigate the relationship between sleep and ASD. However, few studies to date have focused on sleep abnormalities^7^. In earlier work, we reported that individuals with Phelan-McDermid syndrome (PMS), a rare genetic syndrome with high rates of ASD diagnosis, have a sleep phenotype akin to those with ASD^8^. PMS is caused either by loss of the tip of chromosome 22 that includes *SHANK3* or a mutation in the SHANK3 gene, which encodes a neuronal junction protein critical for synaptic function. Mutations in *Shank3* are also often present in idiopathic ASD^9^. Similar to what we observed in patients, we found that adult mice lacking exon 21 of *Shank3* (Shank3^ΔC^) slept less than controls and took longer to fall asleep^8^. Shank3^ΔC^ mice also displayed lower levels of electro-encephalographic (EEG) slow wave (i.e., ‘delta’) activity at baseline showing their sleep was also not of the same quality. Non-rapid eye movement (NREM) sleep delta power dynamics in response to sleep loss are proposed to be a marker of homeostatic sleep pressure. However, Shank3^ΔC^ mice show no differences in NREM sleep delta power dynamics in response to sleep deprivation (SD), suggesting they have no problems accumulating sleep pressure.

In this study, we characterize sleep ontogenesis in Shank3^ΔC^ mice with a view to capture early developmental sleep patterns. To this end, we executed longitudinal sleep recordings starting immediately after weaning into young adulthood. We discovered that mutant mice sleep less overall than wild-type (WT) controls analogous to observations in high-risk infants and toddlers with ASD that go on to be diagnosed. Despite sleeping less overall, Shank3^ΔC^ mice show a significantly increased amount of rapid-eye movement sleep (REM) in early life. We also find that Shank3^ΔC^ mice fail to reduce sleep latency in response to sleep loss as they get older, a physiological response that develops in WT mice between 24 and 30 days of life. These data identify a developmental time window at which early intervention capable of normalizing aberrant sleep patterns could provide therapeutic benefit. Last, we identify features of the EEG spectra that may have potential as an early biomarker of sleep problems in ASD. Overall, our study emphasizes the importance of examining developmental trajectories to understand sleep problems associated with ASD.

## MATERIALS AND METHODS

Animals: Shank3^ΔC^ mice previously characterized by Kouser et al.^10^ on a C57Bl/6 background and available through the Jackson laboratories were bred as previously described^8^, and housed at 24 ± 1°C on a 12:12 hour light: dark cycle with food and water ad libitum. All experimental procedures were approved by the Institutional Care and Use Committee of Washington State University and conducted in accordance with National Research Council guidelines and regulations for experiments in live animals.

Surgical procedures: At postnatal (P) 18 days old, male mice (n=10 Shank3^ΔC^ and n=10 WT littermates) were weaned from their dams and placed under isoflurane anesthesia and stereotaxically implanted with four EEG and two electromyographic (EMG) electrodes as previously described^8^. Briefly, four stainless steel wire loop electrodes were placed bilaterally over frontal (2) and parietal (2) cortices, and EMG electrodes were inserted bilaterally into the nuchal muscles. Adult mice (n=10, approximately 90 days old) were also implanted with four stainless steel screw electrodes (BC-002MP188, Bellcan International Corp, Hialeah, FL) as described above. Bilateral frontal electrode placement in young and adult mice was centered in the frontal skull plates, and bilateral parietal electrodes were placed centrally in the parietal skull plates (exact coordinates at this age vary depending on skull size. To prevent damage to implants, instrumented mice were housed individually from surgery to the completion of final recordings. Mice were allowed a minimum of 3 days of recovery from surgery before habituation to the recording environment. This study is an extension of our previous work in adult male mice^8^, therefore we limited the scope of the current study to males. ASD is four times more prevalent in males than females, therefore phenotypic characterization in animal models of ASD is usually done first in males.

Sleep Recordings: Sleep recordings were conducted in male mice that were 23-60 days old using a longitudinal design. Three days after surgery (P21), mice were connected to a lightweight, flexible tether and allowed two days to habituate to the recording environment. At 23 days old, mice underwent 24 hours undisturbed baseline EEG and EMG recording beginning at light onset (hour 1). The following day (P24), mice were sleep deprived for 3 hour (hours 1-3) via gentle handling starting at light onset as previously described^8^. Mice were allowed 21 hours of recovery sleep (hours 4-12 of the light period and hours 13-24 of the dark period). A total of four 48 hour recordings were repeated when mice were 29-30 days old, 44-45 days old, and 59-60 days old, respectively. Figure 1 outlines the experimental design. Independently, Two 48 hour sleep recordings in a separate cohort of adult (approximately 90 day old) mice were conducted with each animal receiving a single 3 hour or a single 5 hour SD session, spaced 5 days apart.

**Figure 1.**
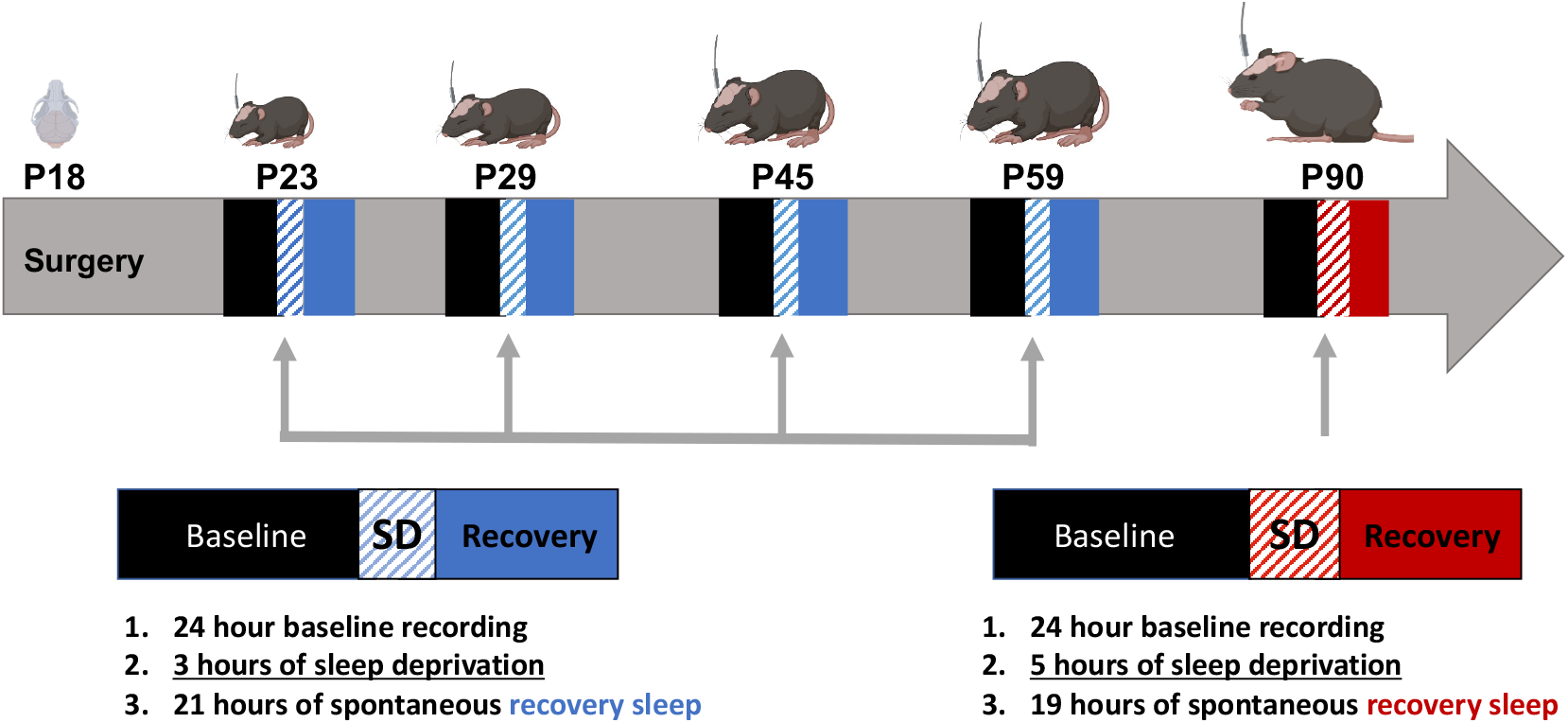
Schematic timeline of experimental procedures. At postnatal day 18 (P18) animals were weaned and surgically implanted for EEG and EMG recordings. Sleep was recorded starting at the following ages: P23, P29, P44 and P59. Animals were recorded for 24 hours of baseline, followed by 3 or 5 hours of sleep deprivation and then 21 hours of spontaneous recovery sleep (48 hours total). Animals from an independent cohort were recorded starting at P90 for 48 hours.

EEG/EMG data acquisition and analysis: EEG and EMG data in animals 23-60 days old were recorded from frontal cortical electrodes (referenced to parietal electrodes) collected with Grass 7 polygraph hardware (Natus Medical Incorporated, Pleasanton, CA) via a light-weight, counterbalanced cable, amplified, and digitized at 256 Hz using VitalRecorder acquisition software (SleepSign for Animal, Kissei Comtec Co., LTD, Nagano, Japan), with band pass filters set at 0.5-30 Hz and notch filtering at 60 Hz. EEG and EMG data in animals 90 days old were collected with Intan RHD2000 Interface using INTAN recording hardware (16-channel RHD USB Recording System, Intan Technologies, Los Angeles CA). EEG and EMG data were recorded from frontal electrodes (referenced to parietal electrodes) at 1 kilo-samples per second with hardware amplification cutoff at 0.01 Hz, lower and upper bandwidths at 0.1 Hz and 200 Hz, and notch filtering at 60 Hz.

Sleep data processing: Recordings were down sampled to 250 Hz using Matlab and exported for manual scoring of sleep states via SleepSign for Animal as previously described^8^. State scoring and data analysis was blinded and randomized. Sleep states and wakefulness were determined by visual inspection of the EEG waveform and EMG activity, and vigilance states were assigned in 4 second increments (epochs). NREM bouts were defined as 7 or more consecutive epochs. REM bouts were defined as 4 or more consecutive epochs. Latency to NREM sleep after SD was defined as time elapsed from release to recovery sleep to the first bout of NREM sleep. The EEG was subjected to fast Fourier transform (FFT) resulting in a power spectrum from 0-20 Hz (P23-P60) or 0-50 Hz (P90) with 0.5 Hz bins. Twelve-hour light period spectra were generated as previously described^8^, from 0.5 Hz spectral bins expressed as a percentage of the sum of total state-specific EEG power (0-20 Hz or 0-50 Hz respectively). NREM delta power (0.5-4 Hz) at baseline was normalized relative to the average NREM power delta band from the last 4 hours of baseline light period (hours 9-12). NREM delta power is dynamic over the course of the day and varies depending on sleep pressure. At the end of the light phase (resting phase for rodents) sleep pressure is minimal and therefore more representative of baseline spectral properties. NREM delta power following SD was normalized relative to baseline NREM delta. Wake theta power (6-9.5 Hz) at baseline was normalized relative to average total power in wake over the 24 hours of baseline. Wake theta power following SD was normalized relative to baseline wake theta. EEG epochs containing artifacts, and recordings with excessive EEG artifacts were excluded from spectral analysis.

Data plotting and statistical Analysis: Statistics were conducted using SPSS for Windows (IBM Corporation Armonk, NY) and RStudio (v. 1.3.1056, RStudio, Boston, MA) as previously described^8^. Non-continuous time-in-state, bout, and latency data are plotted as individual points with a gray bar indicating the group mean. Hourly time-in-state data are presented as means ± standard error of the mean (SEM). Spectra are displayed as smooth curves with 95% confidence intervals, as generated using Generalized Additive Models using the R package mgcv (v.1.8-31). Although our experimental design was longitudinal some animals were excluded from the analysis due to low quality recordings arising from behavioral abnormalities such as excessive repetitive movements or extensive periods of artifact or signal loss at one of the vigilance states. Animals that fell outside ± two standard deviations from the group mean were considered outliers and excluded from analysis. Different animals were excluded at different times. The number of animals excluded per time-point based on data-quality per the specified above criteria is as follows: P23/P24, 3WT and 2 mutants; P29/P30, 3 WT and 2 mutants; P44/P45, 2 WT; P59/P60 2 WT and 2 mutants.

The exclusion of a different set of animals at each time point precluded us from using a repeated measures ANOVA for testing, therefore we implemented a two-way ANOVA with main effects of genotype and age using SPSS for all discrete data. Additional testing across genotype or age was performed post-hoc if main effect of age or genotype were significant. Because we did not detect a significant interaction between age and genotype, post-hoc testing was performed exclusively across genotype within the same age group or across age within genotype. T-tests were used for post-hoc comparisons in all cases, except for hourly time-in-state data, for which 12 hour repeated measures-ANOVAs were used. Significance threshold was set at p ≤ 0.05.

For spectral analysis statistically significant differences were defined by non-overlap of 95% confidence intervals.

## RESULTS

### Developing Shank3^ΔC^ mice sleep less and show altered diurnal/nocturnal distribution of sleep/wake

To understand how the Shank 3 mutation impacts sleep architecture, it is important to first understand normal sleep development. Previous studies established that WT mice by postnatal day 21 exhibit a diurnal/nocturnal (circadian) activity pattern and three distinct states based on EEG: wake, non-rapid eye movement (NREM) sleep and rapid eye movement (REM) sleep^11,12^. How sleep changes after the third postnatal week once the animals are separated from their dams and siblings and into adulthood is not known. The developmental trajectory of sleep in genetic animal models of ASD remains unexplored. To define basal characteristics of postnatal sleep, starting at the third postnatal week, we performed 24-hour EEG and EMG recordings from WT and Shank3^ΔC^ mice under undisturbed (baseline) conditions (at P23, P29, P44, and P59). Even though time asleep in Shank3^ΔC^ mutants is relatively increased during the normal inactive phase (light period), mutants sleep less overall than WT mice at all time points (**Table 1**). In WT animals, we found that the distribution of total sleep time differed significantly between P23 and P29 mice (combined time in NREM and REM) across the light (LP, hours 1 – 12) and dark (DP, hours 13 – 24) periods, measured as the ratio of time asleep in the light period versus the dark period (**Table 1_supplement 1**). These data suggest that diurnal/nocturnal organization of sleep/wake continues to develop between P23 and P29. Although sleep consolidation in the light period in WT mice is still evolving between P23 and P29, Shank3^ΔC^ mice at P23 show the adult ratio of diurnal/nocturnal sleep distribution (**Table 1_supplement 1**).

**Table 1:**
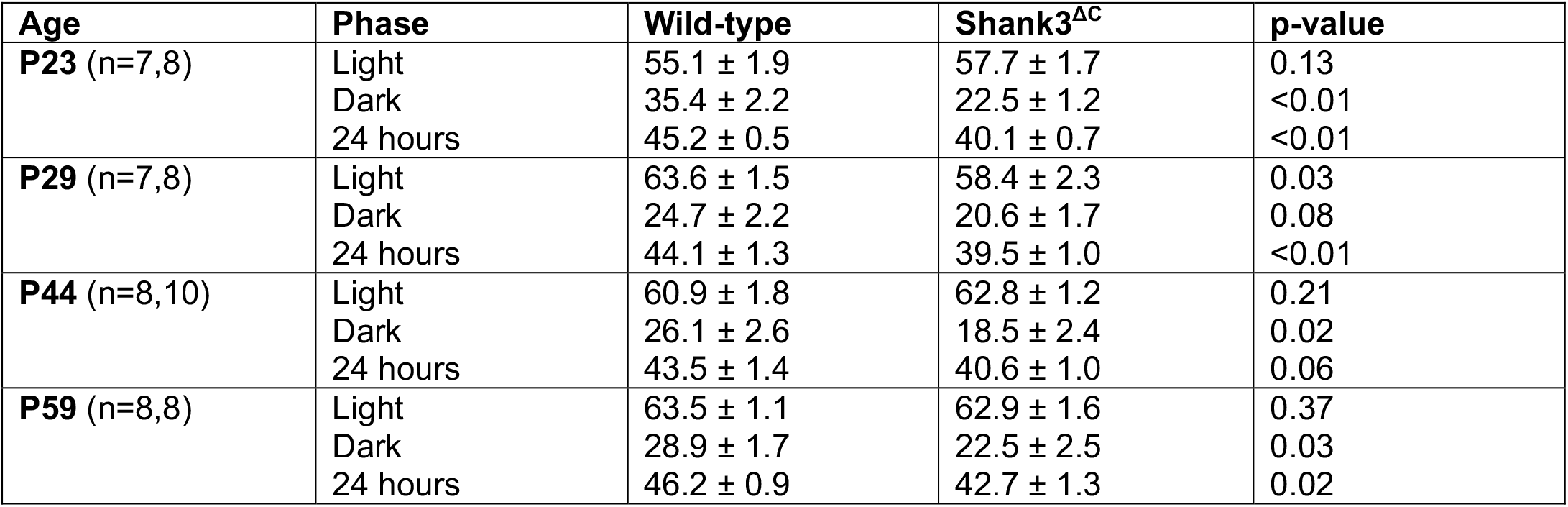
Shank3^ΔC^ mice sleep less throughout their lifespan. Average sleep time during 12 hours of light, 12 hours of dark, or across the full day expressed as percent of total recording time at baseline (24 hours). Standard error is also displayed. Sleep was recorded at postnatal days 23, 29, 44, and 59. Postnatal day 23 (P23) (n=7 WT, 8 Shank3^ΔC^), P29 (n=7 WT, 8 Shank3^ΔC^), P44 (n=8 WT, 10 Shank3^ΔC^), P59 (n=8 WT, 8 Shank3^ΔC^) mice. p-values shown for genotype comparisons at each age (unpaired t-tests).

### Shank3^ΔC^ mice have higher amounts of REM sleep early in life

To determine which states are affected by the overall reduction in sleep we examined time awake, in NREM and in REM sleep over 24 hours starting at P23. At this developmental time point, mice have acquired 70% of their maximal brain volume and are equivalent to a 9-month-old infant based on brain size^13^. The role of sleep during this developmental time period is of particular interest given that results from behavioral studies suggest a period of typical development followed by the early postnatal onset of ASD in the latter part of the first year or early second year of life in human infants^14^.

Similar to what we previously reported in adulthood, increased wakefulness (reduced sleep) in Shank3^ΔC^ mice is most pronounced during the dark period at P23 (p= 0.04; **Figure 2A, Figure 2-supplement 1)**. Time spent in NREM is significantly reduced in Shank3^ΔC^ mice in both light and dark periods across development (P23: LP p<0.01, DP p<0.01; P29: LP p<0.01, DP p=0.04; P44: LP p=0.59, DP p=0.05; P59: LP p=0.04, DP p=0.03; **Figure 2B, Figure 2-supplement 1**). Bouts of NREM sleep are also shorter across development in Shank3^ΔC^ mice, particularly in the dark period (**Figure 2-supplement 2**). Through postnatal development, and relative to WT mice, Shank3^ΔC^ mice show higher REM sleep time from P23 to P59 during the light period (P23 p<0.01; P29 p=0.04; P59 p=0.04; **Figure 2C, Figure 2 – supplement 1**). Increased REM sleep at P23 and P29 in Shank3^ΔC^ mice is driven by an increase in entries into REM during the light period (P23 p<0.01, P29 p=0.03; **Figure 2 – supplement 2**). Together, these findings suggest that state-specific sleep differences in Shank3^ΔC^ mice are developmentally regulated and emerge early post-weaning. Example traces for wake, NREM, and REM sleep across all ages for both mutants and WT can be found in **Figure 2-supplement 3**. Although we detect a significant effect of both age and phenotype across all time-points, particularly during the dark period, the interaction between genotype and age is not quite significant at p<0.05 (**Figure 2-supplement 4)**. This could be due to lack of sufficient power to detect interactions or because during adolescence the effect of the genotype is not as large as during early ages or in adults.

**Figure 2:**
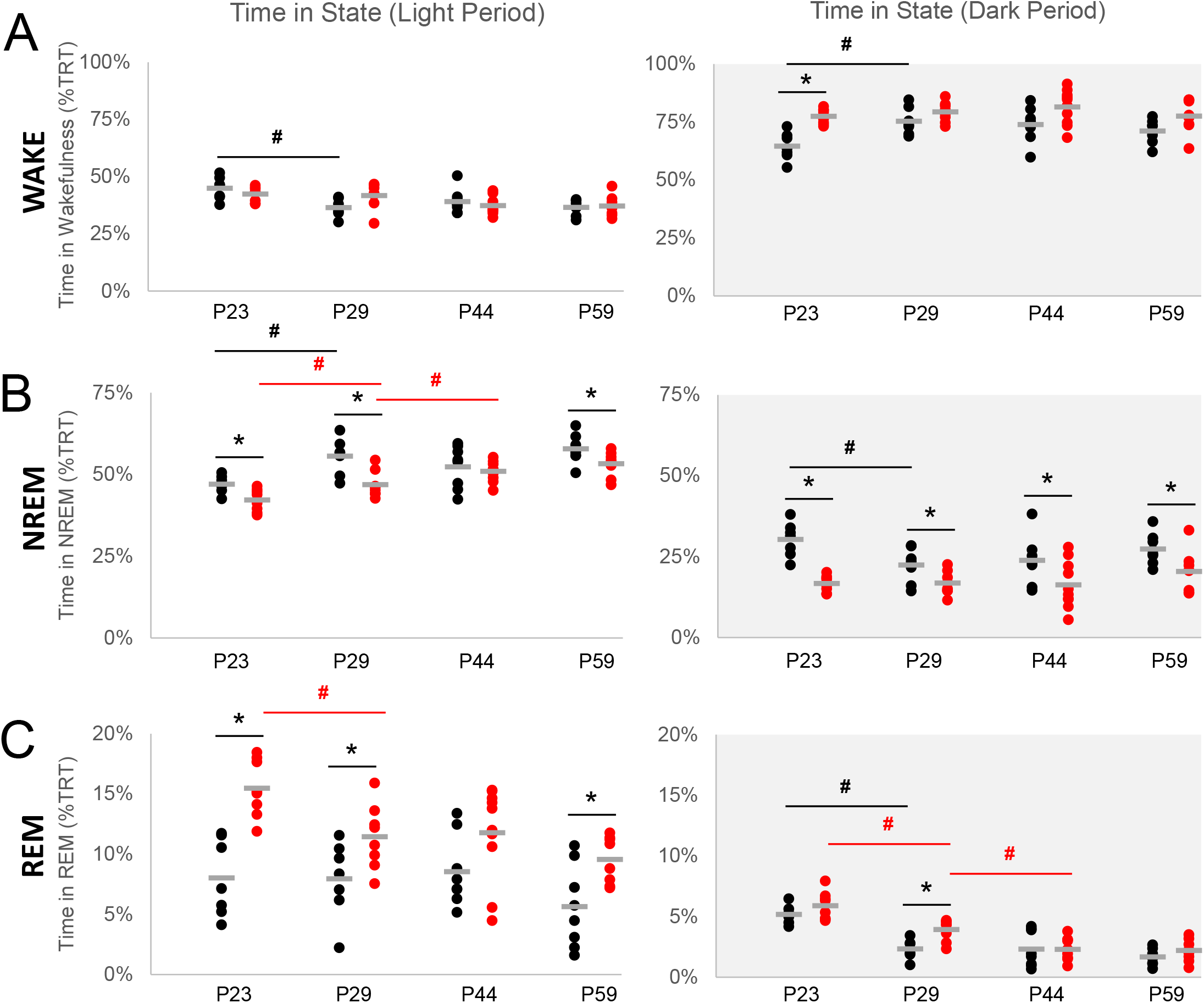
Shank3^ΔC^ mice have increased REM sleep at P23 despite sleeping less. Time (expressed as percent of total recorded time, TRT) in wakefulness (A), NREM sleep (B), and REM sleep (C) during baseline 12 h light (white) and 12 h dark (gray) periods. Sleep was recorded at P23 (n=7 WT, 8 Shank3^ΔC^), P29 (n=7 WT, 8 Shank3^ΔC^), P44 (n=8 WT, 10 Shank3^ΔC^), P59 (n=8 WT, 8 Shank3^ΔC^) mice. Wild-type data is shown in black, Shank3^ΔC^ is shown in red. The average value for each group is shown in grey. * denotes p-values <0.05 obtained from t-test within genotype performed post-hoc after repeated measures ANOVAs within genotype were found significant. # denotes p-values <0.05 obtained from t-test performed post-hoc after two-way ANOVAs (genotype x age) were found significant for genotype or age. No significant interaction between age and genotype was detected (Figure 2-figure supplement 4).

### Spectral power in all brain states changes across development differentially in Shank3^ΔC^ and WT mice

Figure 3. shows the results of spectral analysis across all ages in both mutant and WT mice in wake, NREM and REM sleep. Statistical significance to determine differences across ages within genotype was defined as lack of overlap between 95% confidence intervals. Spectral properties of the rodent cortical EEG waveform are developmentally regulated in a state-specific manner^11,12,15^, and our data in WT animals supports this observation (**Figure 3**). In adult Shank3^ΔC^ mice, we reported that EEG slow-wave delta (0.5-4 Hz) activity in NREM sleep is reduced^8^. Here, we demonstrate that this feature results from a progressive loss of NREM delta activity throughout postnatal development (**Figure 3B, Figure 3-supplement 1, Figure 3-supplement 2, Figure 3-supplement 3**). In WT mice, NREM delta activity in the light period is relatively stable across the same age range, showing only a small reduction between P23 and P29. However, although the power in the theta frequency range (6-9.5 Hz) is gradually reduced through P23 to P59 stages in WT in both the wake and REM spectra, this trend is not present in Shank3^ΔC^ mice, in which the spectral power at this frequency remains the same for all time points (**Figure 3A and 3C**). We also found spectral differences during REM sleep in Shank3^ΔC^ mice, specifically a rightward shift in the peak of theta (6-9.5 Hz) frequency activity and a reduction in overall delta activity (**Figure 3C-supplement 1 and supplement 2**).

### Reduced latency to sleep following sleep loss is a developmentally acquired response that is absent in Shank3^ΔC^ mice

We previously reported that adult Shank3^ΔC^ mice show an increased latency to fall asleep after SD. To better understand how this may develop, we characterized the homeostatic response to SD at P24, P30, P45 and P60 (3 hours, starting at lights on) and recorded changes in sleep/wake architecture and EEGs during the remaining 21 hours. Our results show that at P24 Shank3^ΔC^ mice show no difference in latency to fall asleep following SD relative to WT. However, at P30, Shank3^ΔC^ mice display an increased latency to NREM sleep in Shank3^ΔC^ relative to WT littermates (p=0.04 (**Figure 4A**)). At later time points (P45 and P60), we do not detect differences in latency to sleep across genotypes (**Figure 4-supplement 1)**. This is due to differences in the WT response to SD across ages, because Shank3^ΔC^ mice display the same latency to fall asleep regardless of age (**Figure 4-supplement 2**). Changes in latency to fall asleep following SD as mice get older can have several alternative explanations. First, mice may develop a better ability to stay awake following SD as they get older (SD experiments in adults are traditionally 5-6 hours instead of 3 hours in juveniles). Second, differences in the response to SD may develop during adolescence. Last, there could be long-term effects of being isolated since the electrode placement, that could lead to sleep irregularities. To test the effect of SD length on latency to fall asleep in adult animals, we compared adult P90 animals after 3-hours or 5-hours of SD (**Figure 4-supplement 3**). We show that 3-hours of SD are sufficient to observe the decrease in latency to NREM sleep in adult WT. Interestingly, the difference in latency to fall asleep between adult mutants and WT is larger following 3 hours of SD than 5 hours of SD, suggesting that problems falling asleep in Shank3^ΔC^ mice may be independent from homeostatic sleep pressure. An increase in NREM EEG delta power (0.5-4 Hz) following SD is a commonly used marker of homeostatic sleep pressure; a process that can be detected by the third postnatal week in rodents^11,16,17^. Following 3 hours of SD, we found that WT mice at P24 and at P30 accumulate and discharge NREM delta power in response to SD in the same way as adults^8^ (**Figure 4B, Figure 4-supplement 1, Figure 4-supplement 3**). Theta activity in wakefulness has also been suggested to increase with sleep pressure^18^, although its emergence developmentally has not been examined. We observed genotype differences in theta power at P24 but not at P30 (**Figure 4C**) or later time points (**Figure 4-Figure supplement 1**). Therefore, there does not seem to be a correlation between either NREM delta or wake theta accumulation and difficulties falling asleep in the mutants.

**Figure 3:**
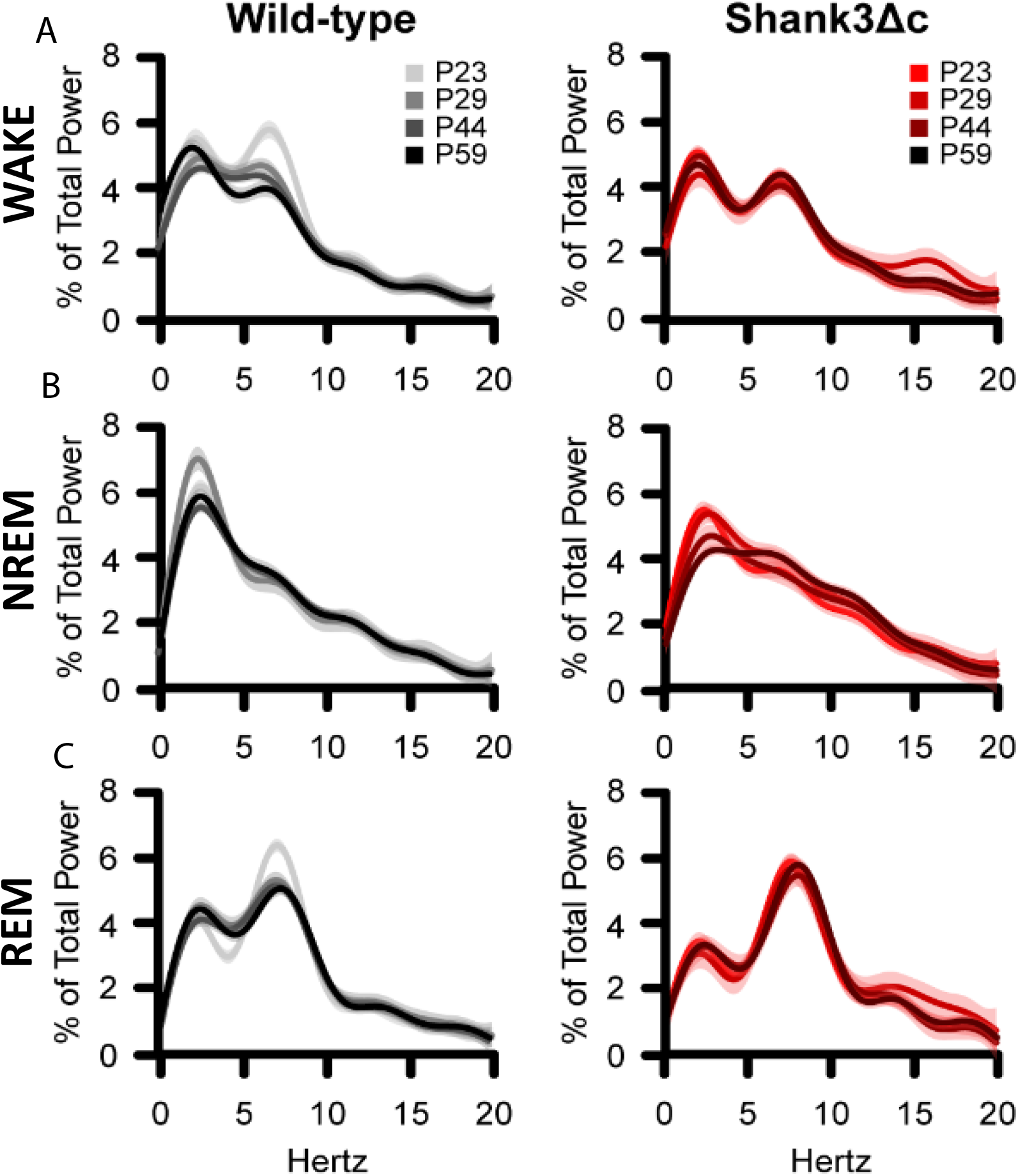
Spectral power changes across development in WT and Shank3^ΔC^ mice. Fourier transformed (FFT) EEG spectral power during the 12 hours of the light period. The rows represent wakefulness (A), NREM sleep (B), and REM sleep (C). EEG spectral power in the light period was normalized as a percentage of total state-specific EEG power in wild-type (left) or Shank3^ΔC^ mice at P23 (n=7 WT, 8 Shank3^ΔC^), P29 (n=7 WT, 8 Shank3^ΔC^), P44 (n=8 WT, 10 Shank3^ΔC^), P59 (n=8 WT, 8 Shank3^ΔC^). Spectra are graphed as smooth lines in grayscale for WT and shades of red for Shank3^ΔC^. 95% confidence intervals are displayed around each spectrum, light gray for WT, and light red for Shank3^ΔC^.

**Figure 4:**
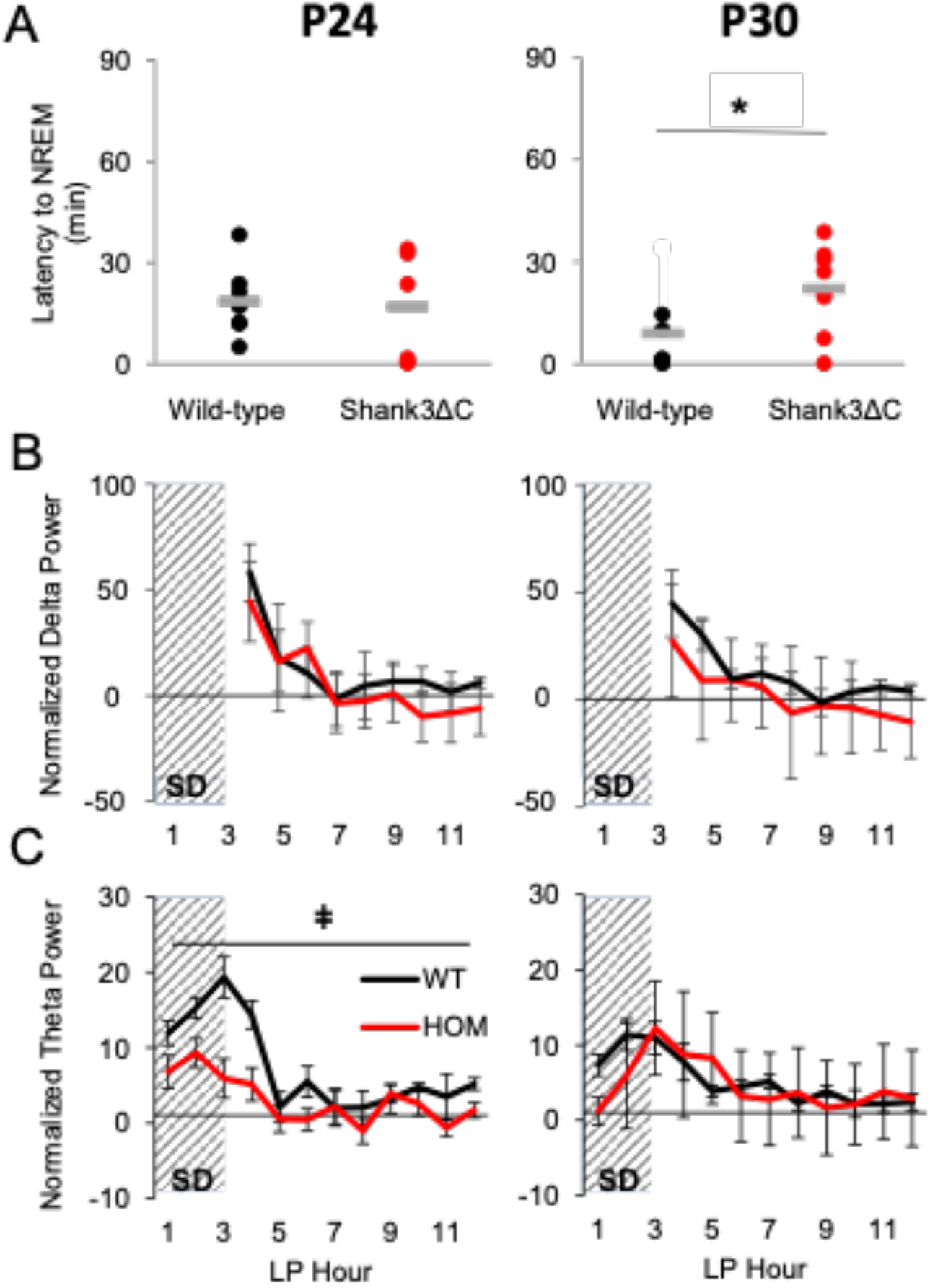
Shank3^ΔC^ mice fail to decrease sleep latency in response to sleep need at P30. A. Latency to the first bout of NREM following 3 hours of SD. P24 (n=7 WT, 8 Shank3^ΔC^), P30 (n=7 WT, 8 Shank3^ΔC^) mice. B. Normalized NREM delta (0.5-4 Hz) power during recovery sleep after 3 hours of SD during the light period (LP) relative to NREM delta power at baseline. C. Normalized Wake theta (6-9.5 Hz) power during 3 hours of SD and subsequent recovery sleep in the light period (LP) relative to Wake theta power at baseline. Repeated measures ANOVA p-values p<0.05 are indicated for genotype differences (*). Wild-type data is shown in black, Shank3^ΔC^ is shown in red, SD period is indicated by crosshatching.

## DISCUSSION

Sleep patterns in children with ASD diverge from typical development early in life, but little is known about the underlying causes. To begin to address this question, we present the first longitudinal trajectory study of postnatal sleep development in the Shank3^ΔC^ ASD mouse model. Our studies highlight that several features of normal sleep are still maturing between P23 and P30 in mice. At 23 days of life, mice have acquired 70% of their maximal brain volume and are equivalent to a 9-month-old infant based on brain size. At P30, mice have reached 80% of their maximal brain volume and are equivalent to an 18 month-old toddler^13^. We find that normal sleep at P23 occurs at a higher proportion in the active phase (night for mice) and is less consolidated (the bouts are shorter). The diurnal-nocturnal distribution of sleep and activity is still maturing between P23 and P29 in WT animals. Shank3^ΔC^ mice at P23 already sleep most of the time in the light phase, like WT mice at P29 do, despite sleeping less overall. This suggests a precocious development of nocturnal/diurnal sleep organization in the mutants.

Shank3^ΔC^ mice show developmental delay in other sleep features. The homeostatic response to sleep loss in young WT mice is different from that of adulthood; at P24, mice take almost 3 times as long to fall asleep following SD than they do at P30. Thus, less consolidated sleep and taking longer to fall asleep are normal features of sleep early in life. Despite sleeping less throughout their lives, Shank3^ΔC^ mice have larger amounts of REM sleep when young, especially at P23. In human brain development, the proportion of REM relative to NREM sleep is greater earlier and dramatically declines upon maturation^19^. Thus, larger amounts of REM suggest that the brain in Shank3^ΔC^ mice is in a more immature state relative to typically developing siblings. Consolidation of NREM sleep into longer bouts is also a normal feature of brain development. As expected from a more immature brain state, Shank3^ΔC^ mice show shorter NREM bouts. The increased REM activity in mutants may arise from an inability to sustain NREM sleep for longer periods of time, i.e., a failure to consolidate sleep. This in turn could underlie another common feature of the ASD sleep phenotype: sleep fragmentation.

EEG spectral analyses show that WT animals overall display more dynamic changes than mutants over time, especially during wake and REM sleep. A characteristic feature of the wake and REM sleep WT EEG spectra at P23 is the presence of a more prominent theta peak. Theta rhythms are mostly hippocampal, and therefore, reduction in this power band at P30 could be explained by an increase in cortical thickness as the animals age. The fact that Shank3^ΔC^ show a change in theta during wake or REM sleep over time may therefore indicate cortical overgrowth at P23, consistent with the early timing of brain overgrowth observed in infants and toddlers that go on to be diagnosed with ASD^20^. More studies will be needed to verify this hypothesis. Nonetheless our data suggest that a reduction of theta power in wake in infancy could serve as an early biomarker of ASD. Interestingly, Shank3^ΔC^ mice also show a reduction in power on the delta frequency range in NREM sleep as they age, suggesting a progressive deterioration in connectivity of the network that underlies slow-wave oscillations in NREM sleep which can explain why sleep problems are reported to worsen over time in ASD.

Taking longer to fall asleep, one of the more salient aspects of the Shank3^ΔC^ adult phenotype, is also a defining characteristic of the clinical ASD sleep phenotype. Latency to sleep onset can only be reliably measured following SD, to make sure that all animals are under comparable sleep pressure. Our study shows that this delay in sleep onset in Shank3^ΔC^ arises from a failure to develop a proper response to sleep loss between 24 and 30 days of age. This deficit arises in the absence of problems accumulating sleep pressure (sleepiness), at least as measured by an increase in delta power in response to SD. In other words, it is a failure of being able to fall asleep quickly despite being sleepy, or reminiscent of being ‘over-tired’. Our studies indicate that this response is normal at young ages. At P24, WT animals take a longer time to fall asleep following sleep loss, but gradually develop the ability to respond to sleep loss by falling asleep faster. In contrast, Shank3^ΔC^ mice display the same latency to fall asleep at all ages. In other words, they maintain an ‘infant’ response to SD throughout their lives. Given that Shank3^ΔC^ mice display the same homeostatic delta response to SD as WT at each age, the absence of genotype differences in sleep latency between P45 and P60 reflect increases in sleep latency in WT animals. This change in WT animals could be driven by a normal increase in latency to fall asleep during adolescence, or due to stress from prolonged social isolation inherent to instrumentation. Regardless of the cause, Shank3^ΔC^ mice seem unable to adjust how long it takes to fall asleep in response to SD as they develop. Our findings regarding the sleep homeostatic response parallel the immature features of baseline sleep we described above and indicate a mis-regulation of normal sleep development in Shank3^ΔC^ mice.

Although the Shank3^ΔC^ sleep phenotype can be considered immature, it may not necessarily arise from a delay in maturation and may in contrast arise from certain aspects of the sleep cycle maturing too early or too fast. For example, Shank3^ΔC^ mutants show a more mature diurnal/nocturnal distribution of sleep/wake at P23. The notion that an early maturation of sleep/wake distribution may explain the eventual failure to develop a proper response to sleep loss may seem counterintuitive. However, it is well supported by our current understanding of sleep regulation. The two-process model of sleep regulation states that when and how much we sleep is determined by the interaction of two processes: the circadian clock and the sleep homeostat^21^. The circadian clock determines the timing of activity during the 24-hour day and is thought to promote arousal during the active phase. The sleep homeostat tracks sleep pressure in response to time spent awake and promotes sleep in response to sleepiness. Both processes mature after the second postnatal week in rodents^22^, although circadian regulation of sleep timing is known to emerge earlier than the ability to track sleep pressure. Based on our observations in WT animals, both regulatory mechanisms may mature independently until the 3^rd^ postnatal week, in which diurnal/nocturnal organization and homeostatic mechanisms may align, then misalign during adolescence, and eventually produce the typical response to sleep loss we observe in adult animals. If the response to circadian input matures too rapidly, one risks too much arousing input from the clock. Excess arousal input from the clock may in turn cause a failure of both processes to crosstalk effectively and explains several features of the Shank3^ΔC^ sleep phenotype, namely: reduced total sleep time, earlier emergence of the diurnal/nocturnal distribution of sleep/wake, shorter NREM sleep bouts, reduced delta power in NREM sleep and delay in sleep onset following SD. A more careful examination of the development of the mechanisms and brain regions that regulate arousal and how they interact with those that promote sleep during the third and fourth postnatal week is warranted to support this hypothesis.

Overall, our results support a reduction of sleep as a core aspect of ASD, while highlighting a period during early life in which the abnormal sleep phenotype fully emerges and may be amenable to intervention. Our results support an excessive arousing input from the circadian clock emerging too early in development as a plausible mechanistic explanation for the observed Shank3^ΔC^ sleep phenotype. Further replication of our findings in other mouse models of ASD is important to dissect which aspects of the sleep phenotype may be specific to the mutation we are modeling. Last, brain development in mice can be substantially different from brain development in humans. Therefore, although rodent studies can be used as broad guidelines, objective longitudinal studies of sleep in both typically developing children and infants at high-risk for ASD are still necessary to determine which of the features we identified in mice apply to ASD in humans.

## CONFLICT OF INTEREST STATEMENT

Financial disclosure: The authors have no financial arrangements or connections to declare.

Non-financial disclosure: The authors have no conflicts of interests to declare.

## AUTHOR CONTRIBUTIONS

All authors had full access to all the data in the study and take responsibility for the integrity of the data. *Conceptualization*, H.S., E.M., and L.P.; *Methodology*, E.M., T.W., K.F., K.S., and H.S.; *Data Curation*, T.W., E.M, K.F., and H.S., *Writing -Original Draft*, L.P.; *Writing - Review & Editing*, H.S, E.M, K.S., and L.P.; *Visualization*, E.M., H.S., and L.P.; *Supervision*, L.P.; *Funding Acquisition*, L.P.

## ACKNOWLEDGEMENTS

We thank Dr. Marcos Frank for valuable discussions. We thank Dr. Paul Worley for providing the Shank3 mutant mouse line. This work was supported by the Justice Equity Diversity and Inclusion (JEDI) award from the Life Science Editors foundation and K01NS104172 from NIH/NINDS to Peixoto L.

## DATA ACCESSIBILITY STATEMENT

The raw EEG data that support the findings of this study are available from the corresponding author upon reasonable request.

## PREPRINT REPOSITORY

This article has been submitted to bioRxiv: https://doi.org/10.1101/2021.03.10.434728

## Notes

**SUPPORT** This work was supported by the Justice Equity Diversity and Inclusion (JEDI) award from the Life Science Editors foundation and K01NS104172 from NIH/NINDS to Peixoto L.

### Competing Interest Statement

The authors have declared no competing interest.

## REFERENCES

1. Maxwell-Horn A, Malow BA. Sleep in Autism. Semin Neurol. 2017;37(4):413–418. doi:10.1055/s-0037-1604353

2. Souders MC, Zavodny S, Eriksen W, et al. Sleep in Children with Autism Spectrum Disorder. Curr Psychiatry Rep. 2017;19(6):34. doi:10.1007/s11920-017-0782-x

3. Hodge D, Carollo TM, Lewin M, Hoffman CD, Sweeney DP. Sleep patterns in children with and without autism spectrum disorders: developmental comparisons. Res Dev Disabil. 2014;35(7):1631–1638. doi:10.1016/j.ridd.2014.03.037

4. MacDuffie KE, Munson J, Greenson J, et al. Sleep Problems and Trajectories of Restricted and Repetitive Behaviors in Children with Neurodevelopmental Disabilities. J Autism Dev Disord. March 2020. doi:10.1007/s10803-020-04438-y

5. MacDuffie KE, Shen MD, Dager SR, et al. Sleep Onset Problems and Subcortical Development in Infants Later Diagnosed With Autism Spectrum Disorder. Am J Psychiatry. 2020;177(6):518–525. doi:10.1176/appi.ajp.2019.19060666

6. Rylaarsdam L, Guemez-Gamboa A. Genetic Causes and Modifiers of Autism Spectrum Disorder. Front Cell Neurosci. 2019;13:385. doi:10.3389/fncel.2019.00385

7. Wintler T, Schoch H, Frank MG, Peixoto L. Sleep, brain development, and autism spectrum disorders: Insights from animal models. J Neurosci Res. March 2020. doi:10.1002/jnr.24619

8. Ingiosi AM, Schoch H, Wintler T, et al. Shank3 modulates sleep and expression of circadian transcription factors. Elife. 2019;8. doi:10.7554/eLife.42819

9. Cochoy DM, Kolevzon A, Kajiwara Y, et al. Phenotypic and functional analysis of SHANK3 stop mutations identified in individuals with ASD and/or ID. Mol Autism. 2015;6:23. doi:10.1186/s13229-015-0020-5

10. Kouser M, Speed HE, Dewey CM, et al. Loss of predominant Shank3 isoforms results in hippocampus-dependent impairments in behavior and synaptic transmission. J Neurosci. 2013;33(47):18448–18468. doi:10.1523/JNEUROSCI.3017-13.2013

11. Nelson AB, Faraguna U, Zoltan JT, Tononi G, Cirelli C. Sleep Patterns and Homeostatic Mechanisms in Adolescent Mice. Brain Sci. 2013;3(1):318–343. doi:10.3390/brainsci3010318

12. Rensing N, Moy B, Friedman JL, Galindo R, Wong M. Longitudinal analysis of developmental changes in electroencephalography patterns and sleep-wake states of the neonatal mouse. Russo E, ed. PLoS ONE. 2018;13(11):e0207031. doi:10.1371/journal.pone.0207031

13. Workman AD, Charvet CJ, Clancy B, Darlington RB, Finlay BL. Modeling Transformations of Neurodevelopmental Sequences across Mammalian Species. J Neurosci. 2013;33(17):7368–7383. doi:10.1523/JNEUROSCI.5746-12.2013

14. Zwaigenbaum L, Bryson S, Rogers T, Roberts W, Brian J, Szatmari P. Behavioral manifestations of autism in the first year of life. Int J Dev Neurosci. 2005;23(2-3):143–152. doi:10.1016/j.ijdevneu.2004.05.001

15. Frank MG, Heller HC. Development of REM and slow wave sleep in the rat. Am J Physiol. 1997;272(6 Pt 2):R1792–1799. doi:10.1152/ajpregu.1997.272.6.R1792

16. Franken P, Chollet D, Tafti M. The homeostatic regulation of sleep need is under genetic control. J Neurosci. 2001;21(8):2610–2621.

17. Frank MG, Morrissette R, Heller HC. Effects of sleep deprivation in neonatal rats. Am J Physiol. 1998;275(1 Pt 2):R148–157.

18. Vassalli A, Franken P. Hypocretin (orexin) is critical in sustaining theta/gamma-rich waking behaviors that drive sleep need. Proc Natl Acad Sci U S A. 2017;114(27):E5464–E5473. doi:10.1073/pnas.1700983114

19. Roffwarg HP, Muzio JN, Dement WC. Ontogenetic development of the human sleep-dream cycle. Science. 1966;152(3722):604–619. doi:10.1126/science.152.3722.604

20. Hazlett HC, Poe M, Gerig G, et al. Early Brain Overgrowth in Autism Associated with an Increase in Cortical Surface Area Before Age 2. Arch Gen Psychiatry. 2011;68(5):467–476. doi:10.1001/archgenpsychiatry.2011.39

21. Borbély AA. A two process model of sleep regulation. Hum Neurobiol. 1982;1(3):195–204.

22. Frank MG, Ruby NF, Heller HC, Franken P. Development of Circadian Sleep Regulation in the Rat: A Longitudinal Study Under Constant Conditions. Sleep. 2016;40(3). doi:10.1093/sleep/zsw077

