## Supplementary materials for "Shank3 influences mammalian sleep development"

**Affiliations:**

**Supplementary materials**

**Table 1_Supplement 1. Shank3$^{\Delta C}$ mice sleep more during the light period.** Ratio of sleep occurring in the light period relative to sleep occurring in the dark period. Sleep was recorded at P23 (n=7 WT, 8 Shank3$^{\Delta C}$), P29 (n=7 WT, 8 Shank3$^{\Delta C}$), P44 (n=8 WT, 10 Shank3$^{\Delta C}$), and P59 (n=8 WT, 8 Shank3$^{\Delta C}$). Wild-type data is shown in black, Shank3$^{\Delta C}$ is shown in red. * denotes p-values <0.05 obtained from t-test within genotype performed post-hoc after repeated measures ANOVAs within genotype were found significant. # denotes p-values <0.05 obtained from t-test performed post-hoc after two-way ANOVAs (genotype x age) were found significant for genotype and age. No significant interaction between age and genotype was detected (Table 1 - supplement 2).

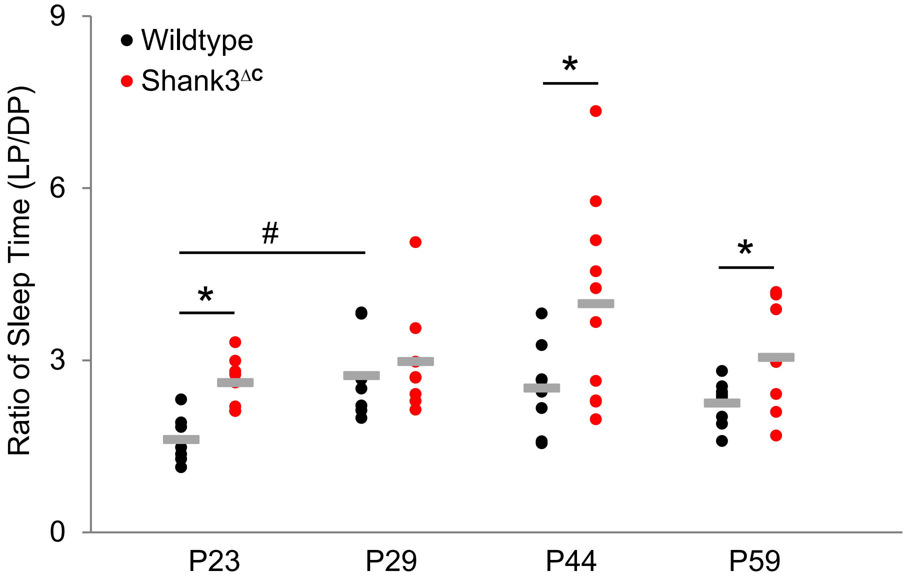

**Table 1- supplement 2. Results of two-way ANOVA (main effect of genotype and age) for the ratio of sleep time (LP/DP).**

Dependent Variable: Ratio of sleep in light period relative to dark period

| Source | Type III Sum of Squares | df | Mean Square | F | Sig. |
|---|---|---|---|---|---|
| Corrected Model | 26.292[a] | 7 | 3.756 | 4.067 | .001 |
| Intercept | 477.245 | 1 | 477.245 | 516.743 | <.001 |
| Age | 12.453 | 3 | 4.151 | 4.494 | .007 |
| Genotype | 10.916 | 1 | 10.916 | 11.819 | .001 |
| Age * Genotype | 1.596 | 3 | .532 | .576 | .633 |
| Error | 51.720 | 56 | .924 | | |
| Total | 579.822 | 64 | | | |
| Corrected Total | 78.012 | 63 | | | |

a. R Squared = .337 (Adjusted R Squared = .254)

**Figure 2 – supplement 1. Shank3$^{\Delta C}$ mice display reduced NREM sleep throughout their lifespan and increased REM sleep when young under baseline conditions.** Time in wakefulness, NREM sleep and REM sleep during baseline 24-hour recordings is shown as a percentage of recording time per hour (average and standard error). Sleep was recorded at P23 (n=7 WT, 8 Shank3$^{\Delta C}$), P29 (n=7 WT, 8 Shank3$^{\Delta C}$), P44 (n=8 WT, 10 Shank3$^{\Delta C}$), P59 (n=8 WT, 8 Shank3$^{\Delta C}$) mice. Statistical significance was determined using repeated measures-ANOVA, main effect of genotype over a 12-hour period. Light period (hours 0-12) and dark period (hours 13-24) were tested separately. * $p<0.05$. WT in black, Shank3$^{\Delta C}$ in red.

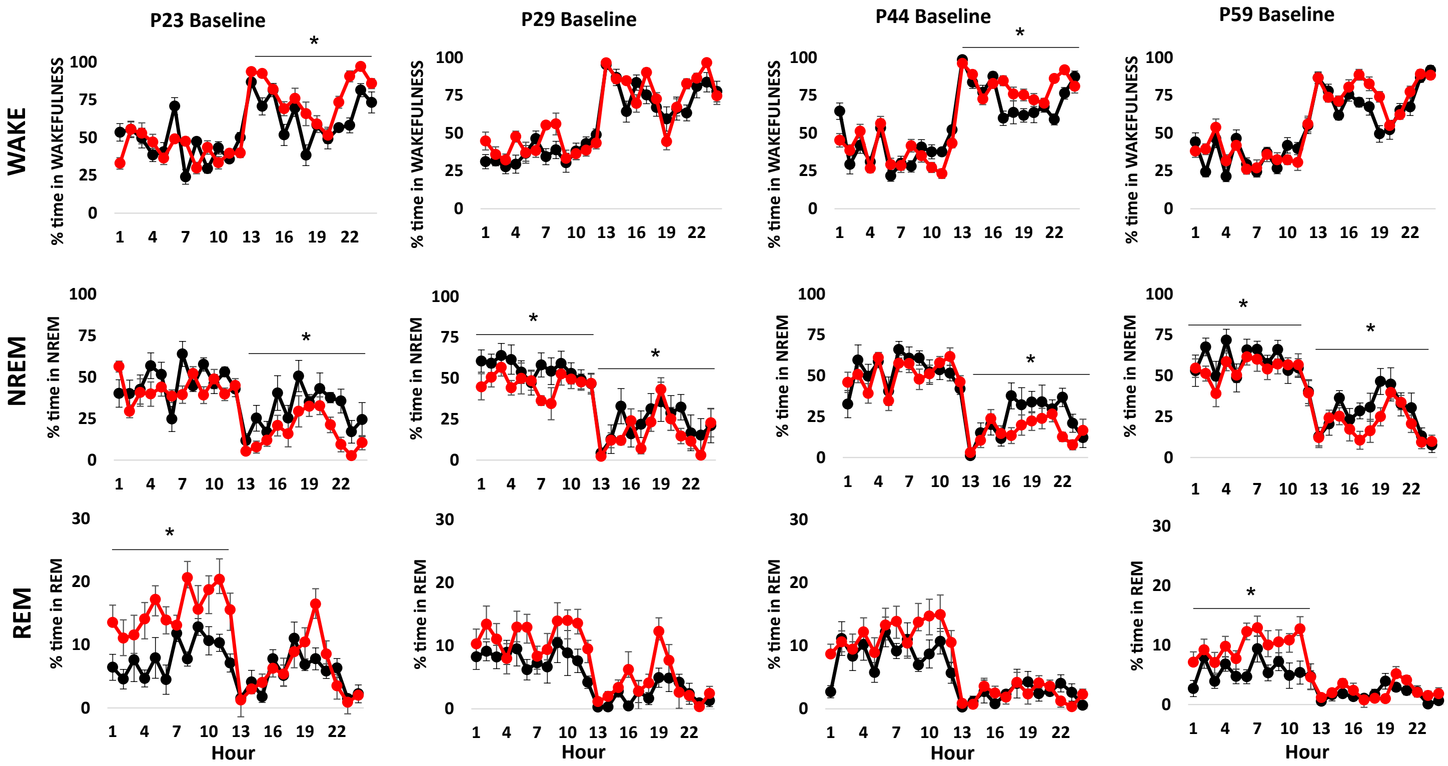

**Figure 2– supplement 2. Age-dependent loss of baseline sleep time in Shank3$^{\Delta C}$ mice is driven by reduced duration of NREM bouts.** The rows represent average number (A-C) and duration (D-F) of bouts of wakefulness (A,D), NREM sleep (B,E), and REM sleep (C,F) during baseline 12 h light (white) and dark (gray) periods. Sleep was recorded at P23 (n=7 WT, 8 Shank3$^{\Delta C}$), P29 (n=7 WT, 8 Shank3$^{\Delta C}$), P44 (n=8 WT, 10 Shank3$^{\Delta C}$), P59 (n=8 WT, 8 Shank3$^{\Delta C}$) mice. * p-value <0.05, repeated measures ANOVA. WT in black, Shank3$^{\Delta C}$ in red.

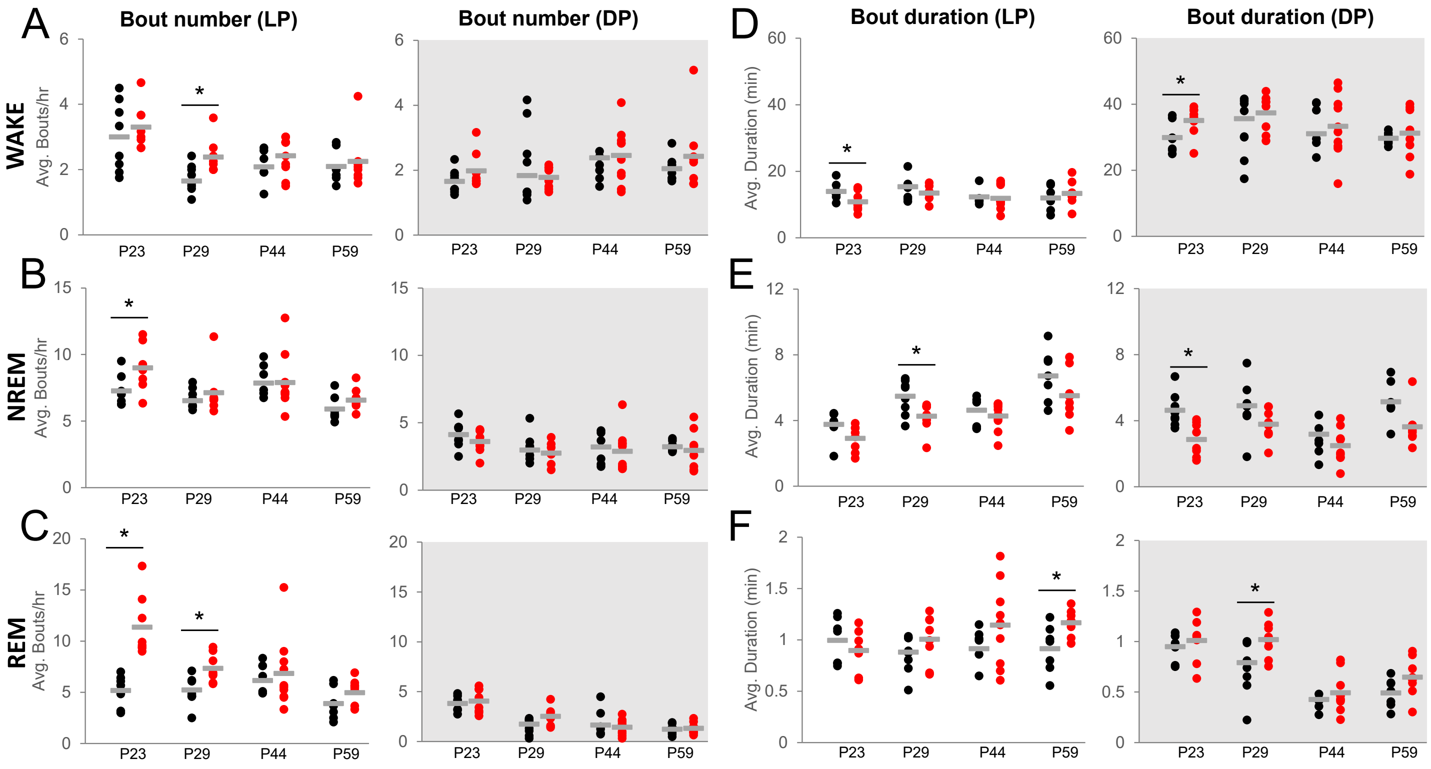

**Figure 2– supplement 3. EEG traces in Shank3$^{\Delta C}$ mice exhibit typical age- and state-specific changes across development.** Example EEG (blue) and EMG (red) traces from Wake, NREM, and REM from one Wild-type and one Shank3$^{\Delta C}$ mouse at P23, P29, P44, P59. Traces are selected from baseline LP recordings and are comprised of two 4-second epochs (denoted by vertical gray lines). Scale and gain are the same across all ages.

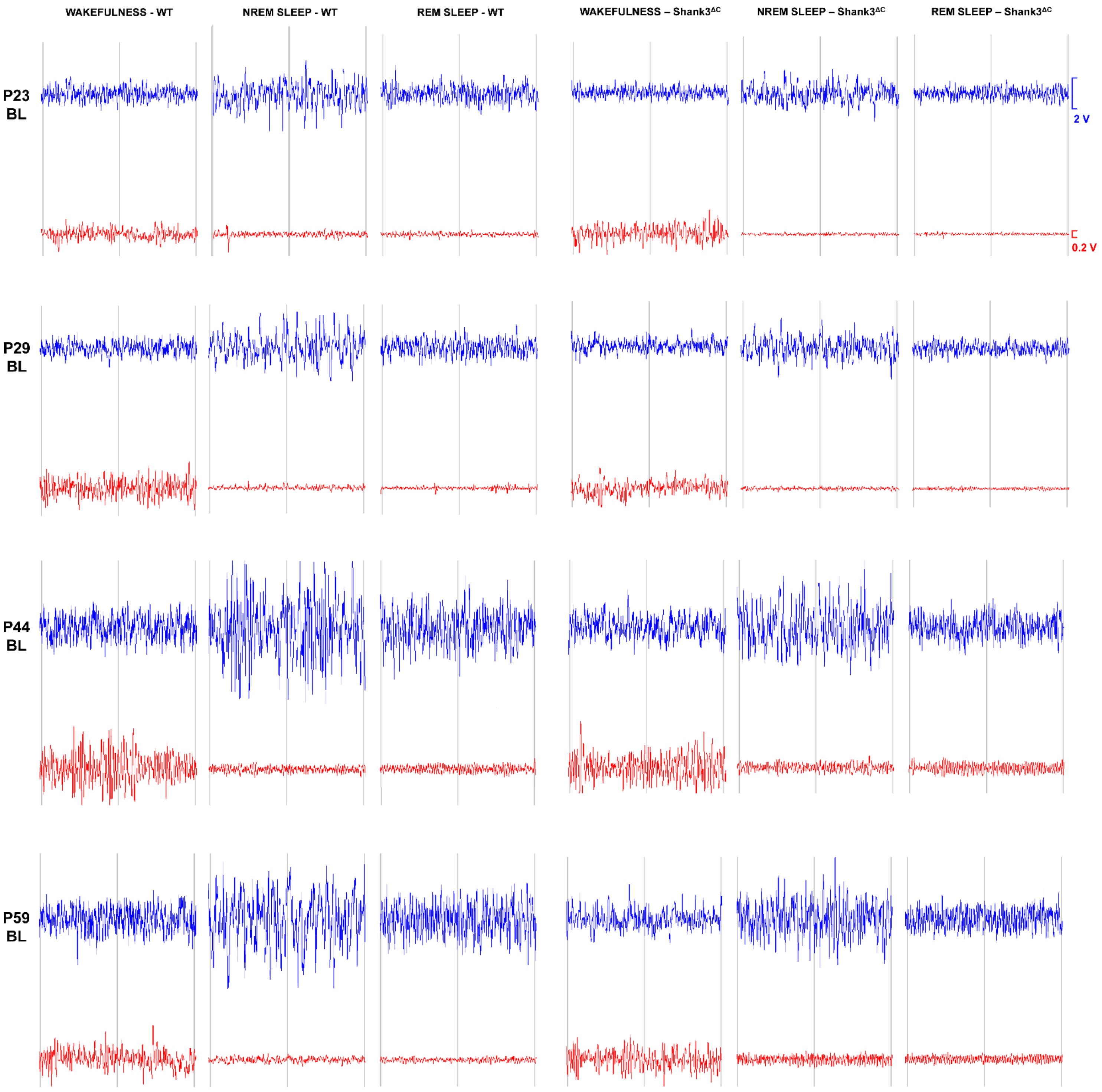

**Figure 2– supplement 4. Results of two-way ANOVAs (main effects of genotype and age) for time in state data.**

Dependent Variable: Wake, Light Period

| Source | Type III Sum of Squares | df | Mean Square | F | Sig. |
|---|---|---|---|---|---|
| Corrected Model | .054[a] | 7 | .008 | 4.087 | .001 |
| Intercept | 9.834 | 1 | 9.834 | 5213.531 | <.001 |
| Age | .040 | 3 | .013 | 7.111 | <.001 |
| Genotype | .000 | 1 | .000 | .113 | .738 |
| Age * Genotype | .014 | 3 | .005 | 2.509 | .068 |
| Error | .106 | 56 | .002 | | |
| Total | 10.045 | 64 | | | |
| Corrected Total | .160 | 63 | | | |

a. R Squared = .338 (Adjusted R Squared = .255)

Dependent Variable: NREM, Light Period

| Source | Type III Sum of Squares | df | Mean Square | F | Sig. |
|---|---|---|---|---|---|
| Corrected Model | .145[a] | 7 | .021 | 11.253 | <.001 |
| Intercept | 16.334 | 1 | 16.334 | 8891.325 | <.001 |
| Age | .095 | 3 | .032 | 17.296 | <.001 |
| Genotype | .038 | 1 | .038 | 20.622 | <.001 |
| Age * Genotype | .011 | 3 | .004 | 1.977 | .128 |
| Error | .103 | 56 | .002 | | |
| Total | 16.749 | 64 | | | |
| Corrected Total | .248 | 63 | | | |

a. R Squared = .584 (Adjusted R Squared = .533)

Dependent Variable: REM, Light Period

| Source | Type III Sum of Squares | df | Mean Square | F | Sig. |
|---|---|---|---|---|---|
| Corrected Model | .051[a] | 7 | .007 | 8.006 | <.001 |
| Intercept | .609 | 1 | .609 | 662.452 | <.001 |
| Age | .014 | 3 | .005 | 4.959 | .004 |
| Genotype | .032 | 1 | .032 | 35.288 | <.001 |
| Age * Genotype | .004 | 3 | .001 | 1.614 | .196 |
| Error | .051 | 56 | .001 | | |
| Total | .733 | 64 | | | |
| Corrected Total | .103 | 63 | | | |

a. R Squared = .500 (Adjusted R Squared = .438)

Dependent Variable: Wake, Dark Period

| Source | Type III Sum of Squares | df | Mean Square | F | Sig. |
|---|---|---|---|---|---|
| Corrected Model | .154[a] | 7 | .022 | 6.053 | <.001 |
| Intercept | 35.690 | 1 | 35.690 | 9801.229 | <.001 |
| Age | .045 | 3 | .015 | 4.112 | .010 |
| Genotype | .094 | 1 | .094 | 25.819 | <.001 |
| Age * Genotype | .016 | 3 | .005 | 1.441 | .240 |
| Error | .204 | 56 | .004 | | |
| Total | 36.776 | 64 | | | |
| Corrected Total | .358 | 63 | | | |

a. R Squared = .431 (Adjusted R Squared = .360)

Dependent Variable: NREM, Dark Period

| Source | Type III Sum of Squares | df | Mean Square | F | Sig. |
|---|---|---|---|---|---|
| Corrected Model | .150[a] | 7 | .021 | 6.709 | <.001 |
| Intercept | 2.981 | 1 | 2.981 | 933.139 | <.001 |
| Age | .023 | 3 | .008 | 2.443 | .074 |
| Genotype | .112 | 1 | .112 | 35.065 | <.001 |
| Age * Genotype | .014 | 3 | .005 | 1.477 | .231 |
| Error | .179 | 56 | .003 | | |
| Total | 3.256 | 64 | | | |
| Corrected Total | .329 | 63 | | | |

a. R Squared = .456 (Adjusted R Squared = .388)

Dependent Variable: REM, Dark Period.

| Source | Type III Sum of Squares | df | Mean Square | F | Sig. |
|---|---|---|---|---|---|
| Corrected Model | .014[a] | 7 | .002 | 20.041 | <.001 |
| Intercept | .066 | 1 | .066 | 680.612 | <.001 |
| Age | .012 | 3 | .004 | 41.652 | <.001 |
| Genotype | .001 | 1 | .001 | 8.147 | .006 |
| Age * Genotype | .001 | 3 | .000 | 1.867 | .146 |
| Error | .005 | 56 | 9.686E-5 | | |
| Total | .084 | 64 | | | |
| Corrected Total | .019 | 63 | | | |

a. R Squared = .715 (Adjusted R Squared = .679)

**Figure 3- supplement 1: Shank3$^{\Delta C}$ mice show progressive reduction in NREM delta activity and a shift in the peak of REM theta activity in the light period.** The rows represent vigilance states of wakefulness (top), NREM sleep (middle), and REM sleep (bottom). EEG spectral power in the light period normalized as a percentage of total state-specific EEG power at P23 (n=7 WT, 8 Shank3$^{\Delta C}$), P29 (n=7 WT, 8 Shank3$^{\Delta C}$), P44 (n=8WT, 10Shank3$^{\Delta C}$), P59 (n=8 WT, 8 Shank3$^{\Delta C}$). Spectra are graphed as smooth lines in black for WT and red for Shank3$^{\Delta C}$. 95% confidence intervals are displayed around each spectrum, light gray for WT, and light red for Shank3$^{\Delta C}$. Frequency in the x-axis is in hertz.

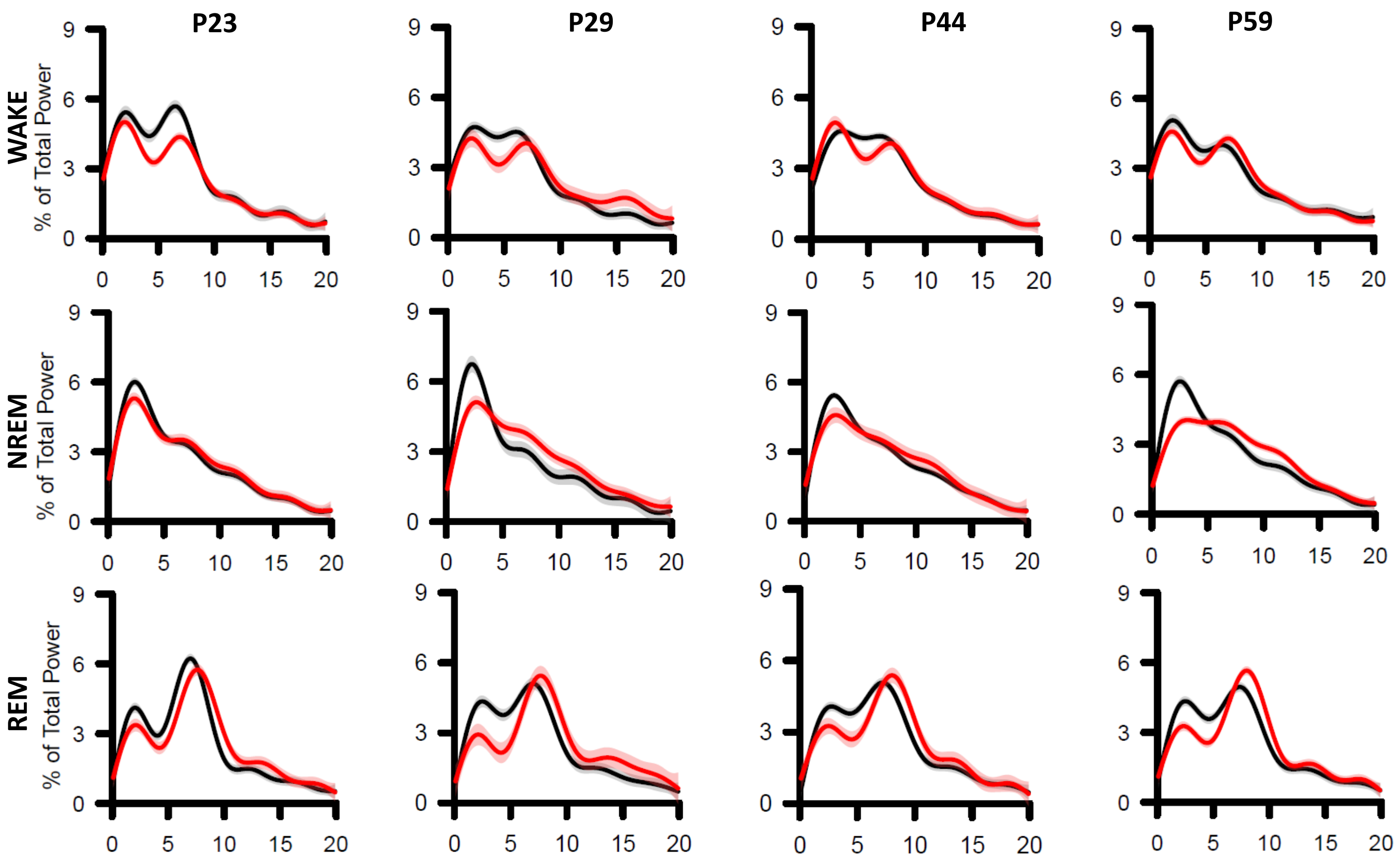

**Figure 3- supplement 2: Age-specific spectral differences in Shank3$^{\Delta C}$ mice are also present in the dark period.** The rows represent vigilance states of wakefulness (top), NREM sleep (middle), and REM sleep (bottom). EEG spectral power in the dark period normalized as a percentage of total state-specific EEG power at P23 (n=7 WT, 8 Shank3$^{\Delta C}$), P29 (n=7 WT, 8 Shank3$^{\Delta C}$), P44 (n=8 WT, 10 Shank3$^{\Delta C}$), P59 (n=8 WT, 8 Shank3$^{\Delta C}$). Spectra are graphed as smooth lines in black for WT and red for Shank3$^{\Delta C}$. 95% confidence intervals are displayed around each spectrum, light gray for WT, and light red for Shank3$^{\Delta C}$. Frequency in the x-axis is in hertz.

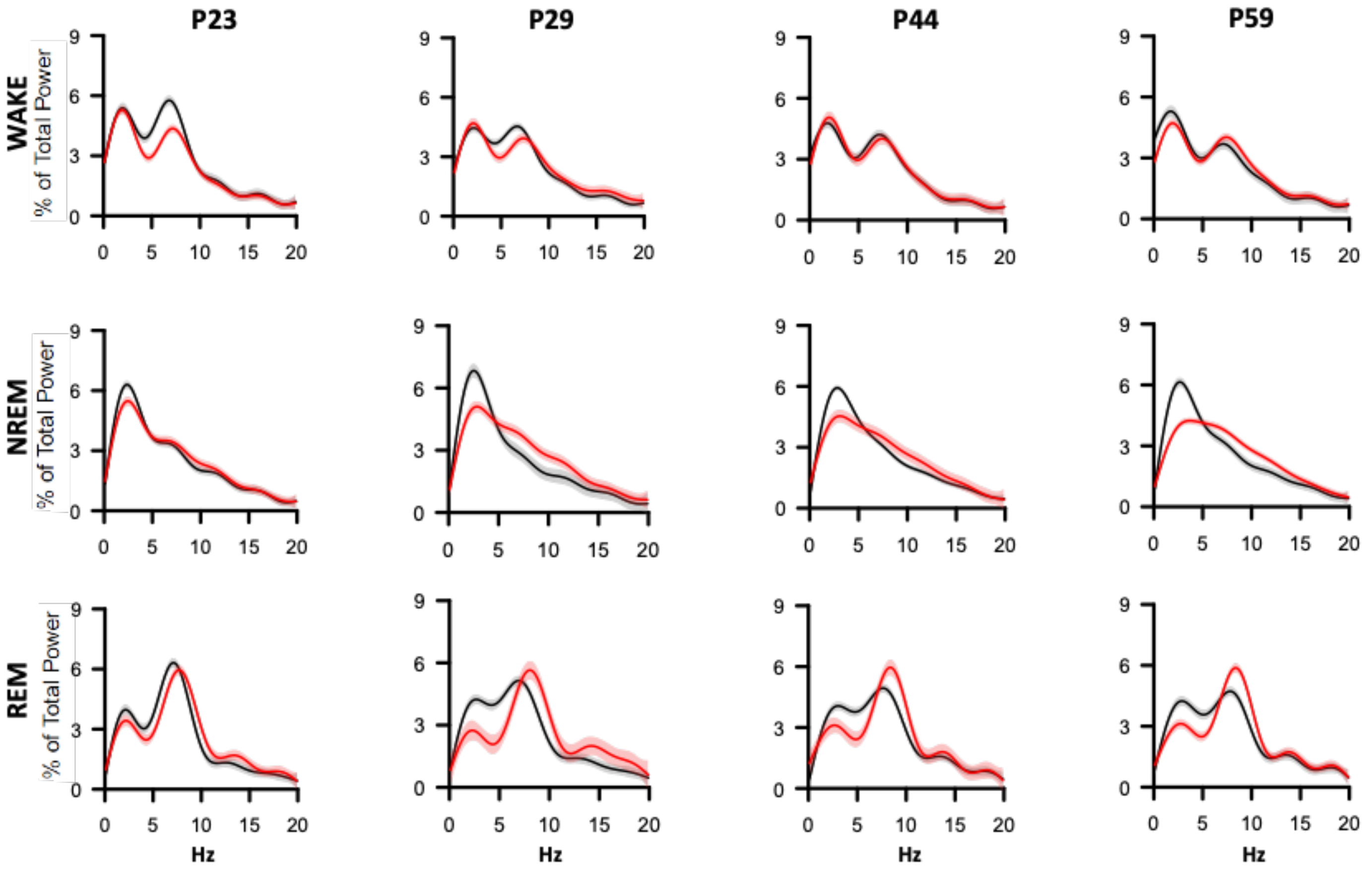

**Figure 3- supplement 3: Shank3$^{\Delta C}$ mice at baseline show reduced NREM delta activity across all ages, and increased wake theta activity at P23.** Normalized delta (0.5-4 Hz) power in baseline NREM sleep (top) and normalized theta (6-9.5 Hz) during wakefulness (bottom). EEG spectral power was normalized as a percentage of total state-specific EEG power at P23 (n=7 WT, 8 Shank3$^{\Delta C}$), P29 (n=7 WT, 8 Shank3$^{\Delta C}$), P44 (n=8 WT, 10 Shank3$^{\Delta C}$), P59 (n=8 WT, 8 Shank3$^{\Delta C}$). * ANOVA p-value<0.05, main effect of genotype is indicated by (*). Wildtype data is shown in black, Shank3$^{\Delta C}$ is shown in red.

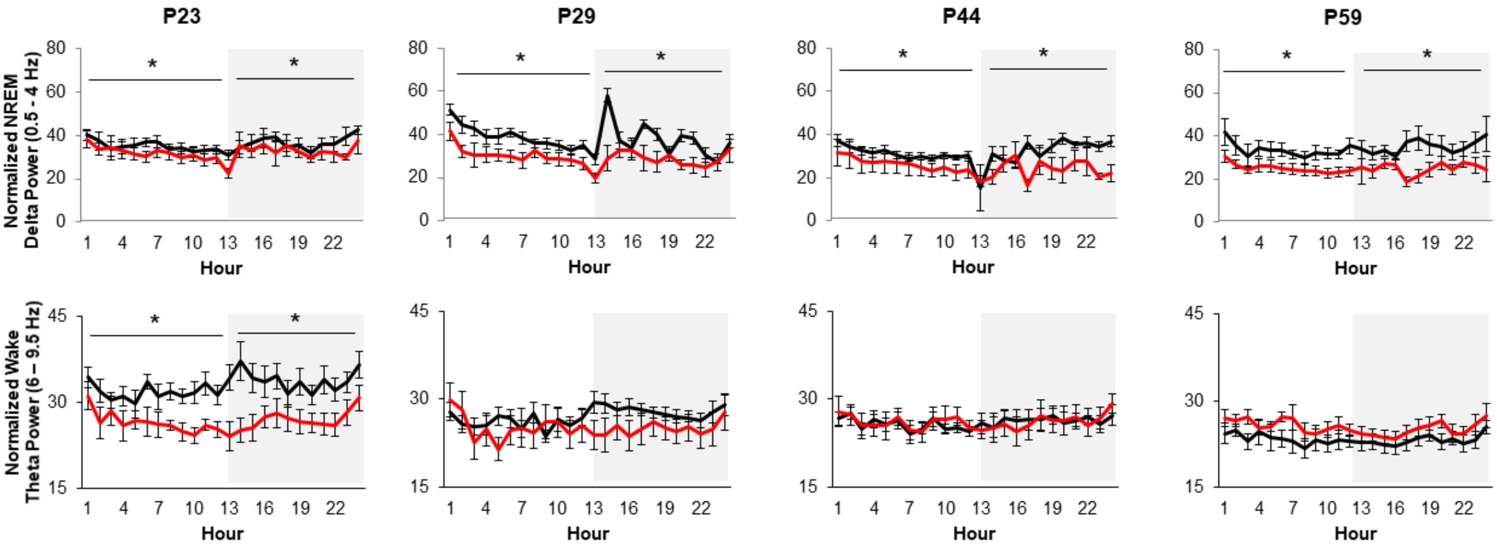

**Figure 4- supplement 1. Sleep latency following SD is unchanged at P45 and P60.** A. Latency to the first bout of NREM sleep following 3 hours of sleep deprivation at P45 (n=8 WT, 10 Shank3$^{\Delta C}$), and P60 (n=8 WT, 8 Shank3$^{\Delta C}$) mice. B. Normalized NREM delta (05-4 Hz) power during recovery sleep after 3 hours of sleep deprivation during the light period (LP) relative to NREM delta power at baseline. C. Normalized WAKE theta (6-9.5 Hz) power during 3 hours of sleep deprivation and subsequent recovery sleep in the light period (LP) relative to WAKE theta power at baseline. Wildtype data is shown in black, Shank3$^{\Delta C}$ is shown in red, SD period is indicated by crosshatching.

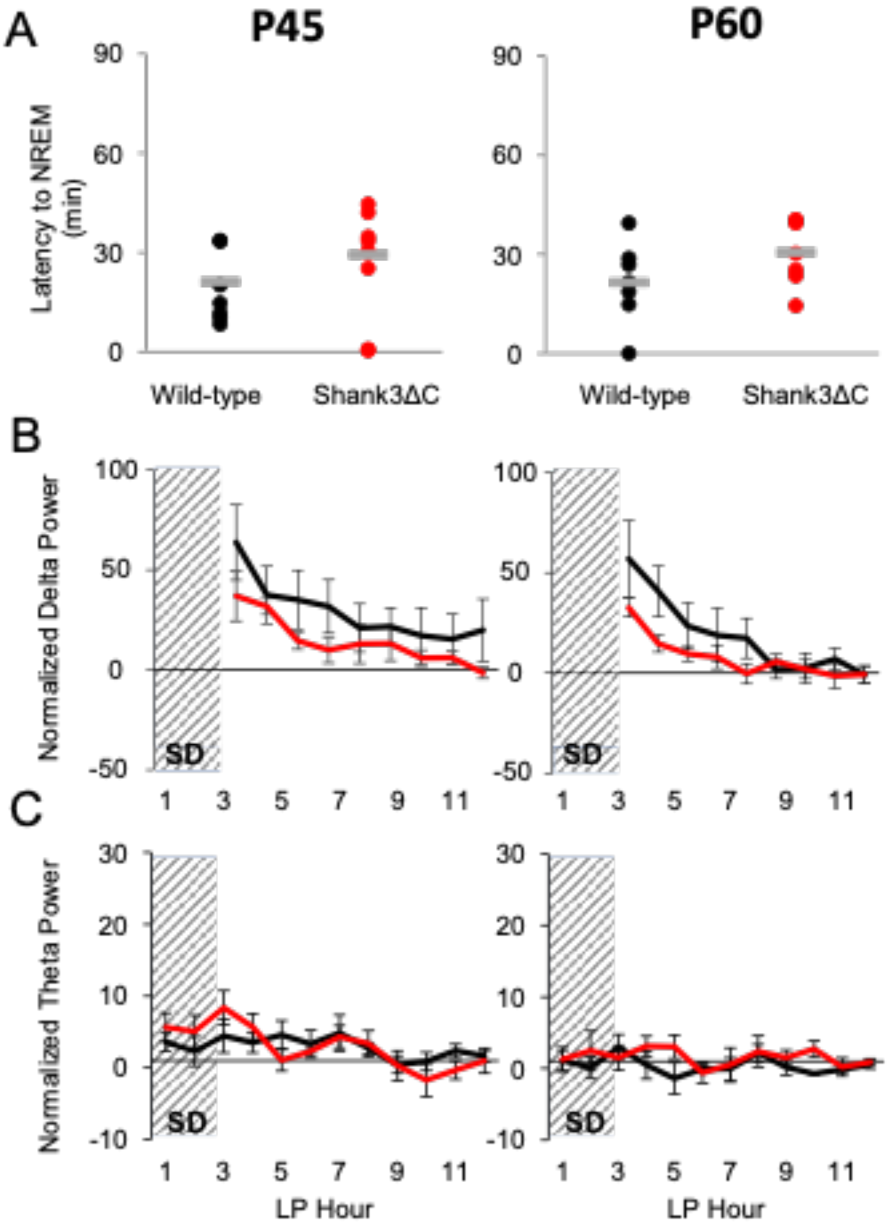

**Figure 4- supplement 2. Latency to fall asleep changes over time in WT but not in Shank3$^{\Delta C}$ mice.** Difference in latency (in minutes) to the first bout of NREM sleep following 3 hours of sleep deprivation across time-points. P24 (n=7 WT, 8 Shank3$^{\Delta C}$), P30 (n=7 WT, 8 Shank3$^{\Delta C}$), P45 (n=8 WT, 10 Shank3$^{\Delta C}$), P60 (n=8 WT, 8 Shank3$^{\Delta C}$), and P90 (n=10 WT, 10 Shank3$^{\Delta C}$) mice. * p-value<0.05, two-tailed unpaired t-tests.

| **Differences in latency to fall asleep after 3 hours of SD across time-points** | | | | |
|---|---|---|---|---|
| *Within-genotype comparisons* | | | | |
| **Wild-type** | | | | |
| | Difference in latency to | Fold Change | p-value (t-test) | number of replicates |
| P24 v. P30 | -14 | 0.239 | 0.02 | n=7 vs n=7 |
| P30 v. P45 | 16 | 4.517 | 0.01 | n=7 vs n=8 |
| P45 v. P60 | 0 | 1.018 | 0.95 | n=8 vs n=8 |
| P60 v. P90 | -14 | 0.357 | 0.01 | n=8 vs n=10 |
| **Shank3$^{\Delta C}$** | | | | |
| | Difference in latency to | Fold Change | p-value (t-test) | Number of replicates |
| P24 v. P30 | 5 | 1.31 | 0.52 | n=8 vs n=8 |
| P30 v. P45 | 7 | 1.323 | 0.37 | n=8 vs n=10 |
| P45 v. P60 | 1 | 1.044 | 0.87 | n=10 vs n=8 |
| P60 v. P90 | -3 | 0.894 | 0.65 | n=8 vs n=10 |

**Figure 4- supplement 3. Sleep latency is increased in adult (P90) mice following either 3 or 5 hours of SD.** A. Normalized NREM delta (0.5-4 Hz) power during recovery sleep after 3 hours of sleep deprivation during the light period (LP) relative to NREM delta power at baseline. B. Latency to the first bout of NREM sleep following 3 hours or 5 hours of sleep deprivation at P90 (n=10 WT, 10 Shank3$^{\Delta C}$). Wildtype data is shown in black, Shank3$^{\Delta C}$ is shown in red, SD period is indicated by crosshatching.

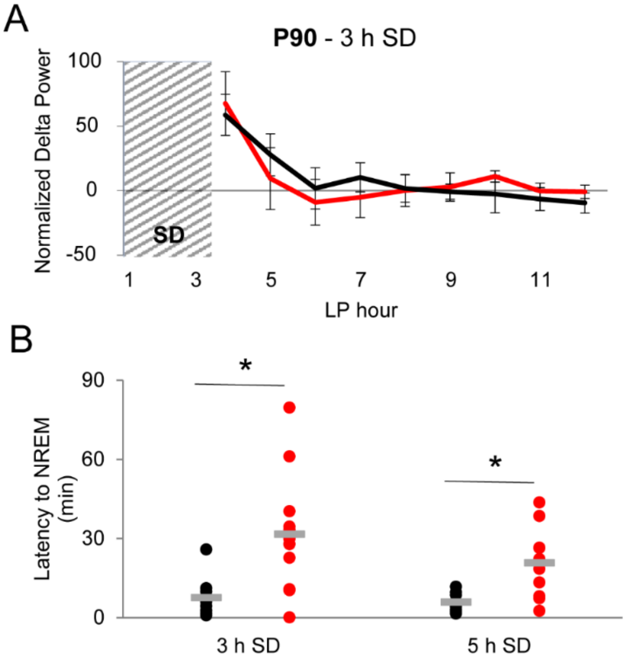
